# easyMF: A Web Platform for Matrix Factorization-based Biological Discovery from Large-scale Transcriptome Data

**DOI:** 10.1101/2020.12.21.405563

**Authors:** Wenlong Ma, Siyuan Chen, Jingjing Zhai, Yuhong Qi, Shang Xie, Minggui Song, Chuang Ma

**Affiliations:** State Key Laboratory of Crop Stress Biology for Arid Areas, Center of Bioinformatics, College of Life Sciences, Northwest A&F University, Shaanxi, Yangling 712100, China; Key Laboratory of Biology and Genetics Improvement of Maize in Arid Area of Northwest Region, Ministry of Agriculture, Northwest A&F University, Shaanxi, Yangling 712100, China

**Keywords:** Galaxy, Integrative analysis, Matrix factorization, Metagene, Transcriptome, RNA Sequencing

## Abstract

With the development of high-throughput experimental technologies, large-scale RNA sequencing (RNA-Seq) data have been and continue to be produced, but have led to challenges in extracting relevant biological knowledge hidden in the produced high-dimensional gene expression matrices. Here, we present easyMF, a user-friendly web platform that aims to facilitate biological discovery from large-scale transcriptome data through matrix factorization (MF). The easyMF platform enables users with little bioinformatics experience to streamline transcriptome analysis from raw reads to gene expression and to decompose expression matrix from thousands of genes to a handful of metagenes. easyMF also offers a series of functional modules for metagene-based exploratory analysis with an emphasis on functional gene discovery. As a modular, containerized and open-source platform, easyMF can be customized to satisfy users’ specific demands and deployed as a web server for broad applications. easyMF is freely available at https://github.com/cma2015/easyMF. We demonstrated the application of easyMF with four case studies using 940 RNA sequencing datasets from maize (*Zea mays* L.).

## Introduction

High-throughput sequencing of RNA (RNA-Seq) is being used in almost all biology and related research laboratories, and has become a key research tool for profiling genome-wide gene expression in various species. The constant improvement of RNA-Seq technologies coupled with sharp decreases in sequencing costs and data generation timelines now enables investigators to perform sequencing-based projects for hundreds of thousands of samples from different cells, tissues, organs, experimental conditions, individuals and species (Cardoso-Moreira et al., 2019; Nelms and Walbot, 2019; One Thousand Plant Transcriptomes, 2019; Sarropoulos et al., 2019; Shulse et al., 2018; Qiu et al., 2020). The large-scale transcriptome sequencing, however, results in considerable challenges for data analysis, as the outputs are naturally represented as high-dimensional gene expression matrices (genes in rows and samples in columns), from which it is difficult to extract new information through traditional gene expression analysis approaches like differential expression analysis and correlation-based statistical analysis.

Machine learning is a branch of artificial intelligence that enables computer algorithms to learn hidden knowledge from Big Data in biology and other sciences (Ma et al., 2014; Mooney and Pejaver, 2018; Cuocolo et al., 2019). Matrix factorization (MF; also known as matrix decomposition) is a class of unsupervised machine learning techniques that can decompose high-dimensional data into low-dimensional structures, while preserving as much information as possible from the original data (Koren et al., 2009). With the development of a variety of computer algorithms, such as principal component analysis (PCA) (Abdi and Williams, 2010), independent component analysis (ICA) (Hyvarinen and Oja, 2000), and non-negative matrix factorization (NMF) (Lee and Seung, 2000), MF is regarded as well suited for large-scale transcriptome data analysis (Stein-O’Brien et al., 2018). MF reduces the gene expression matrix from thousands of genes to a handful of metagenes, each of which can represent a weighted combination of the individual genes. MF can also decompose the gene expression matrix into a product of two low-dimensional matrices: the amplitude matrix (AM; genes in rows and metagenes in columns) and the pattern matrix (PM; metagenes in rows and samples in columns), which have served as the basis for a series of metagene-base applications, such as sample clustering analysis, functional gene discovery, cell type identification, and so on (Stein-O’Brien et al., 2018; Noor et al., 2019; Sompairac et al., 2019; Nguyen and Wang, 2020). Several MF-based pipelines are available, but these tools were designed for specific or limited functionalities (**Table S1**). Moreover, when developing tools for high-throughput sequencing data, ensuring reliability, reproducibility, flexibility and ease of use become a crucial desideratum. Accordingly, the absence of a reliable, reproducible, all-in-one, and easy-to-use platform is to a great extent obstructing MF-based transcriptome analyses for both computational and experimental biologists.

To address this limitation, we here present easyMF, a web platform that facilitates MF-based knowledge discovery from large-scale transcriptome data. The easyMF platform was equipped using the Big-Data-supported Galaxy system with user-friendly graphic user interfaces, allowing researchers with little programming experience to streamline transcriptome analysis from raw reads to gene expression, and to carry out our MF, and metagene-based exploratory analysis. All analysis data, such as inputs, parameters, intermediate results, and outputs, are permanently recorded in the “History” panel of easyMF, making complex MF-based transcriptomic analysis reproducible and amenable to collaborative modes. In addition to the Galaxy system, easyMF was also powered with the advanced Docker packaging technology, making it easy to install and deployable in user-customized hardwire under different operating systems (i.e., Windows, Linux, and Macintosh). With these flexible, interactive, reproducible, and easy-to-use features, we expect easyMF to serve as a valuable tool with broad application potential. We provide examples of the application of easyMF to 940 RNA sequencing datasets of maize (*Zea mays* L.) inbred line B73.

## Results

### Overview of easyMF

The easyMF platform comprises three functional modules, named Matrix Preparation, Matrix Factorization, and Deep Mining (**Figure 1**). Matrix Preparation was designed to prepare a high-quality gene expression matrix for downstream analysis. Matrix Factorization can be used to decompose the gene expression matrix into an AM and PM using three different computer algorithms, i.e., PCA, ICA, and NMF. Deep Mining was designed to perform metagene-based statistical analysis for sample clustering, signature gene identification, functional gene discovery, cell type detection, and pathway activity inference. These functional modules were built with a comprehensive set of functions (**Table S2**), which can be selected by users to customize their own pipelines for satisfying specific needs.

**Figure 1.**
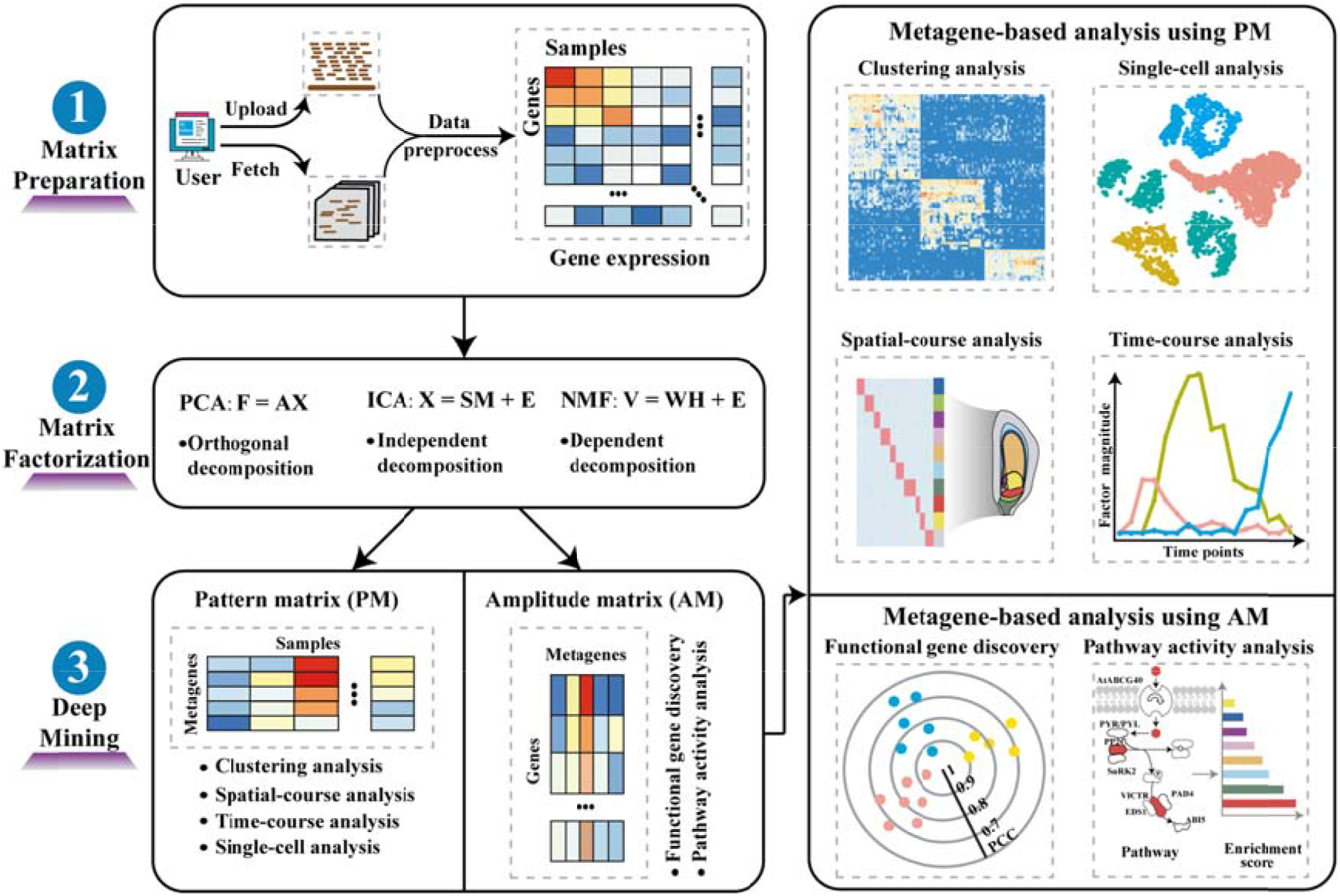
Overview of easyMF.

The easyMF platform is typically started with an input of a gene expression matrix, in which genes are in rows and samples are in columns. The gene expression matrix can also be automatically generated from raw reads using a bioinformatics pipeline (**Figure S1**), which was specially designed for users unfamiliar with RNA-Seq data analysis. After specifying the accession numbers of RNA-Seq datasets from the National Center for Biotechnology Information (NCBI) Gene Expression Omnibus (GEO) and/or Sequence Read Archive (SRA) databases, the customized pipeline can be implemented for a series of RNA-Seq data analyses, including data retrieval, format transferring, quality control, reads mapping, and gene expression quantification. To improve the quality of the gene expression matrix, easyMF removes batch effects from different experiments using the sva function (Leek et al., 2012), filters genes expressed at low levels with user-specified criteria, and removes outlier samples using a sample-based PCA approach (Fehrmann et al., 2015).

The easyMF platform subsequently decomposes the gene expression matrix into a product of the AM and PM with three optional algorithms, namely PCA, ICA and NMF, which calculate metagenes through orthogonal decomposition, independent decomposition and dependent decomposition, respectively. The number of metagenes can be specified by users, or chosen according to optimized parameters: the internal consistency of Cronbach’s α value for PCA (Fehrmann et al., 2015) and the inflection point of the rate of the mean residual decline for ICA and NMF (Gaujoux and Seoighe, 2010). For each metagene, genes with dominant patterns (defined as signature genes) are identified using patternMarkers (Stein-O’Brien et al., 2017) and the Pearson’s correlation coefficient (PCC) algorithm (see **Methods; Figure S2**). The patternMarkers calculates the Euclidean distance between normalized AM coefficients and the 0-1 pattern of metagenes. While the PCC algorithm scores the association between gene expression values and PM coefficients.

The easyMF platform makes use of gene-level relationships in the AM for functional gene discovery (Fehrmann et al., 2015) and pathway activity inference (see **File S1**). This platform also makes use of sample-level relationships in the PM to perform temporal, spatial, and integrated transcriptome analysis. In the current version, easyMF provides six optional algorithms (mclust (Scrucca et al., 2016), apcluster (Bodenhofer et al., 2011), SSE, fpc (Hennig, 2013), vegan (Dixon, 2003), and gap (Maechler et al., 2012)) to cluster samples using PM coefficients. The clusters are visualized in plots, as well as tables, providing a quick overview of the relationships between samples. The easyMF platform can also be used to determine the extent to which genes change over time in response to perturbations (e.g., developmental time), and does so by integrating gene expression values, and gene- and sample-level relationships. It can also be used to identify signature genes dominated at specific compartments of the transcriptomes with spatial resolution in individual tissue samples (spatial transcriptomes), and to identify the type of unknown cells from single-cell RNA-Seq data.

### Application of easyMF to 940 maize RNA-Seq samples

To demonstrate the utility of easyMF, we used it to perform a large-scale analysis of RNA-Seq data from maize B73 samples manually collected from the NCBI GEO and SRA databases (**Table S3**). After a series of data processing steps (see **Methods**), a maize gene expression matrix (denoted as G_1_) of 28,874 protein-coding genes and 940 samples was constructed, in which each gene had FPKM (fragments per kilobase of transcript per million mapped reads) ≥ 1 in at least 15 RNA-Seq samples. As one of the most important sources of food, feed, and biofuel materials, maize seed has been extensively characterized using RNA-Seq technologies to understand its complex gene expression patterns at the genome-wide level. The availability of extensive transcriptomes from 285 seed samples provided us an opportunity to explore the ability of easyMF to be used to attain new knowledge about seeds.

### The easyMF platform is capable of effectively prioritizing seed-related genes

A schematic overview of the application of easyMF to gene prioritization is depicted in **Figure 2A and Figure S3**. easyMF first uses the PCA algorithm to decompose the matrix G_1_ into two matrices, namely amplitude matrix AM_1_ and pattern matrix PM_1_. At a threshold of Cronbach’s *α* > 0.7, easyMF generated 161 metagenes, capturing 96.4% of the variation in gene expression (**Figure S4**). Then, the performance of easyMF in maize functional gene prioritization was extensively evaluated using the leave-one-out cross-validation (LOOCV) strategy on 75 Gene Ontology (GO) terms (**Table S4**), each of which consisted of 5~500 experimentally validated genes, provided by Ensembl Plants (Bolser et al., 2017) (http://plants.ensembl.org) and maize-GAMER (Wimalanathan et al., 2018). For each GO term, we quantified the performance of easyMF using the area under the receiver operating characteristic (ROC) curve (AUC) and the area under the self-ranked curve (AUSR) (for details, see **Methods**). For a comparison, we also tested the recently proposed network-based gene discovery system MaizeNet (Lee et al., 2019) and a random selection strategy using the same LOOCV experiment. The MaizeNet system prioritizes functional genes in maize using a co-functional network inferred from more than 20 distinct types of genomic and proteomic data sets. The random selection process was repeated 100 times by randomly assigning gene identifiers to the score and rank results obtained from PCA in each round of the LOOCV experiments. The mean evaluation results were used for the random selection strategy. For AUC-based and AUSR-based evaluations, easyMF performed much better than random selection, and exhibited comparable or superior prediction performances as compared to the network-based approach MaizeNet (**Figure 2B, C**).

**Figure 2.**
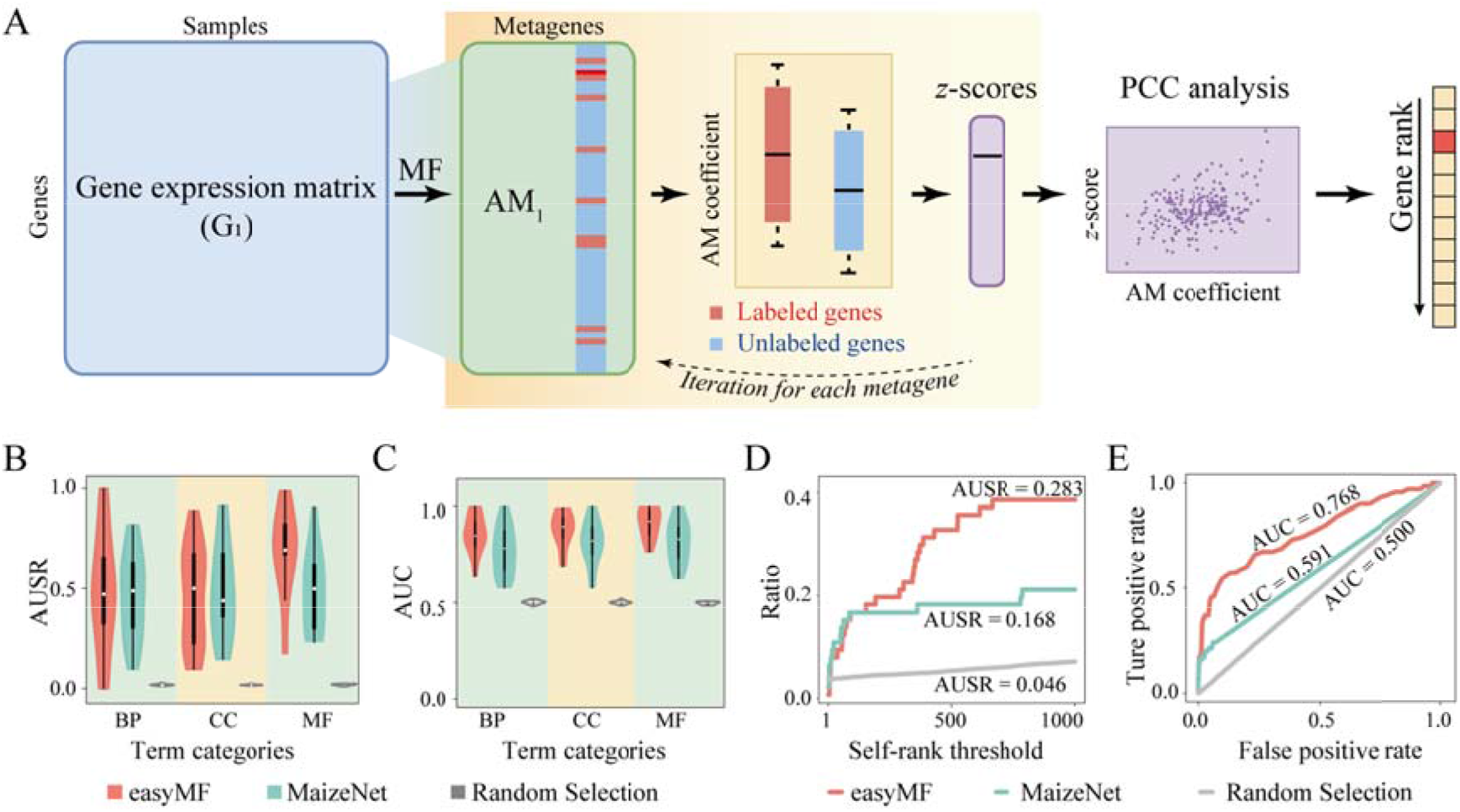
Application of easyMF to gene prioritization. (A) A schematic overview of easyMF in gene prioritization. Performance evaluation of easyMF, MaizeNet and Random Selection approaches in terms of (B) AUSR and (C) AUC. LOOCV experiments were performed using experimentally validated genes from 75 GO terms from Biological Process (BP), Cellular Component (CC), and Molecular Function (MF) domains. The easyMF platform was found to be superior to MaizeNet in the prioritization of 70 seed-related genes in terms of (D) the AUSR, and (E) the AUC.

These encouraging results prompted us to further access the ability of easyMF to prioritize seed-related genes. A manual literature survey was conducted to identify 70 experimentally validated genes functionalized in maize seed development (**Table S5**). The LOOCV experiments on these 70 seed-related genes showed AUSR values of 0.283, 0.168, and 0.046 (**Figure 2D**) and AUC values of 0.768, 0.591, and 0.500 (**Figure 2E**) for easyMF, MaizeNet, and random selection, respectively. Using all of these 70 seed-related genes as input, easyMF generated a prediction model to identify seed-related candidate genes at the genome scale. A detailed literature review showed that four of the top 10 candidates predicted by easyMF have been experimentally validated: *ZmNRP1* (*no-apical-meristem-related protein1*, Zm00001d040189) (Guo et al., 2003; Haun and Springer, 2008; Yi et al., 2019), *ZmMYB127* (*MYB-transcription factor 127*, Zm00001d041935) (Bernardi et al., 2019; Yi et al., 2019), *ZmTAR3* (*tryptophan aminotransferase related3*, Zm00001d037674) (Bernardi et al., 2012; Zhan et al., 2018) and *ZmEREB167* (*AP2-EREBP-transcription factor 167*, Zm00001d032095) (Bernardi et al., 2019; Yi et al., 2019) (**Table S6**).

Overall, these results indicated easyMF to be a reliable and effective platform for prioritizing functional genes through MF-based transcriptome analysis. Lists of seed-related candidate genes prioritized by easyMF and MaizeNet are provided in **Table S6** for the benefit of researchers who in the future may pursue experimental validation.

### The easyMF platform can be used to perform robust sample clustering for facilitating the identification of seed signature genes

We next considered the application of easyMF to sample clustering of a large-scale gene expression matrix. By implementing the NMF algorithm, easyMF decomposed the gene expression matrix G_1_ into two matrices, namely amplitude matrix AM_2_ and pattern matrix PM_2_, and reduced the dimension of G_1_ from 28,874 genes to 11 metagenes (**Figure 3A**). Maize samples can then be analyzed by summarizing gene expression patterns in terms of the coefficients of metagenes (i.e., the relative weights of samples in PM_2_). There were three metagenes (metagene1, metagene7, and metagene10) that had significantly higher coefficients in seed samples than in non-seed samples (**Figure 3B**), indicative of an association between these three metagenes and seed samples. This association was further highlighted by a hierarchical clustering analysis of the PM_2_ (11 metagenes × 940 samples), in which all seed samples were clustered into three subgroups (**Figure 3A**).

**Figure 3.**
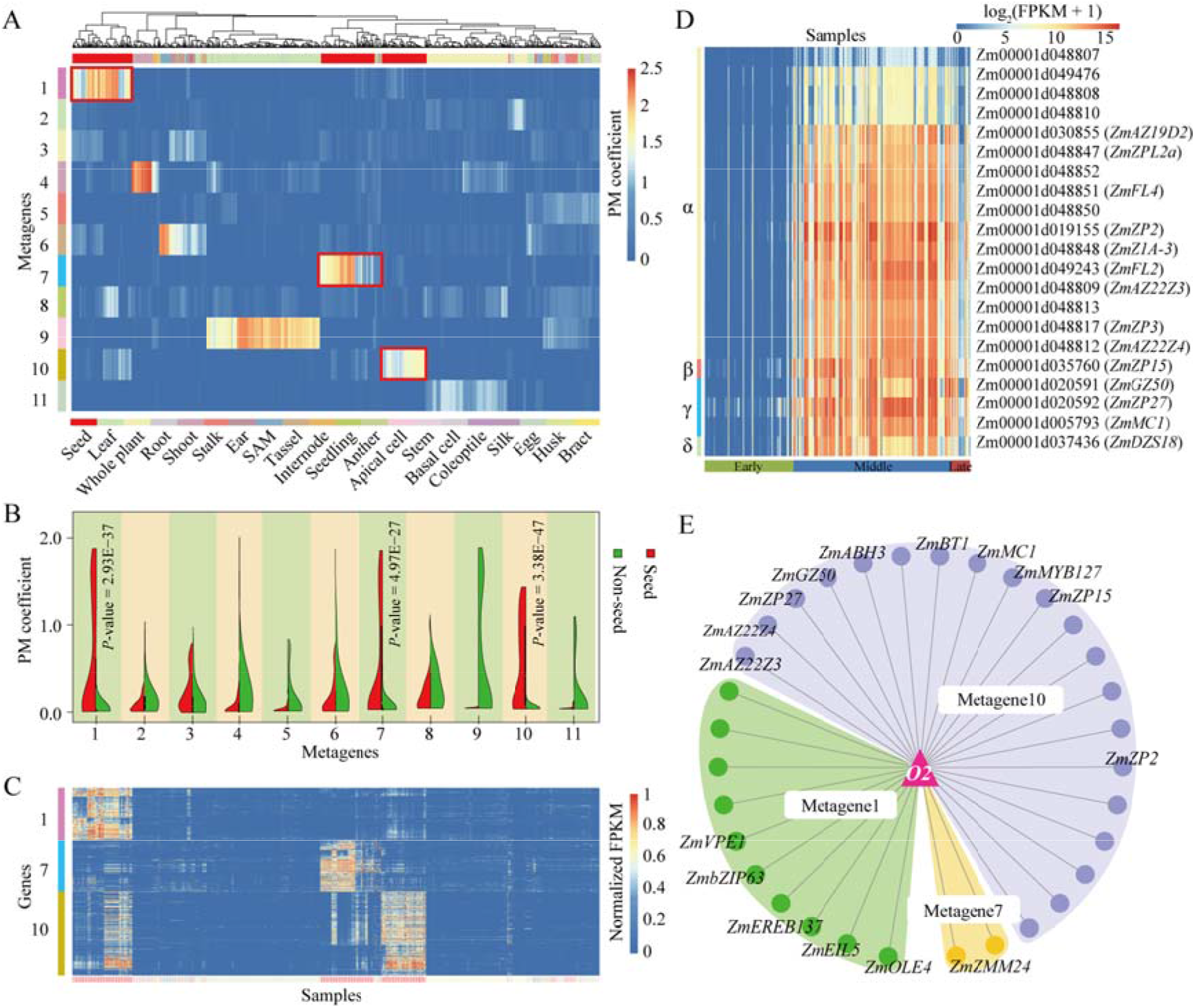
Application of easyMF to clustering analysis. (A) Hierarchical clustering analysis of the pattern matrix PM_2_ decomposed from 940 maize RNA-Seq samples using easyMF. (B) The distribution of coefficients between seed and non-seed samples among 11 metagenes in PM_2_. (C) Expression patterns of seed-related signature genes among 940 maize RNA-Seq samples. (D) Heatmap showing expression levels of 21 zein-encoding genes among 285 seed samples. (E) Regulatory network consisting of *O2* transcription factor and seed-related signature genes. Thirty-two seed-related signature genes were identified as *O2*-modulated and/or -bound target genes using ChIP-Seq data.

Based on these three seed-related metagenes, we identified 774 signature genes (metagene1: 216, metagene7: 213, and metagene10: 345) by using patternMarkers and the PCC algorithm (**Supplemental Table S7**). Most (95.99%) of these 774 genes were specially expressed in seed samples (**Figure 3C**), with this expression pattern determined using the Tau method (Kryuchkova-Mostacci and Robinson-Rechavi, 2017), by which a tissue specificity score higher than 0.7 was measured. Several of these signature genes have been experimentally associated with maize seed development, including *ZmABI3* (*ABSCISIC ACID INSENSITIVE3*; Zm00001d001838) (Ma et al., 2019), *ZmDE18* (*defective18*; Zm00001d023718) (Bernardi et al., 2012), *ZmNAC130* (*NAC-transcription factor 130*, Zm00001d008403) (Zhang et al., 2019), *ZmZAG2* (*Zea AGAMOUS homolog2*, Zm00001d041781) (Schmidt et al., 1993), *ZmSBT2* (*subtilisin2*, Zm00001d006669) (Lopez et al., 2017), and endosperm-specific transcription factors (TFs) *Opaque2* (*O2*; Zm00001d018971) (Schmidt et al., 1990) and *Opaque11* (*O11*; Zm00001d003677) (Feng et al., 2018). In maize kernel, the major chemical component is starch, which provides ~70% of the kernel weight (Flint-Garcia et al., 2009). Of the above 774 signature genes, several were starch-related genes. One representative example is *ZmBT1* (*brittle endosperm1*; Zm00001d015746), a mutant of which severely reduces starch content (Shannon et al., 1998). Maize kernels also contain several types of storage proteins, most of which are zeins (Tsai, 1979). There were 21 zein-encoding genes identified as seed-related signature genes with obviously high expression levels in both the middle and late phases of seed development from 10 days after pollination (DAP), covering four different types of subfamilies including α-, β-, γ-, and δ-zeins (**Figure 3D**). Gene ontology (GO) enrichment analysis revealed several seed-related signature genes associated with embryonic development, including two genes encoding seed maturation proteins (Zm00001d026037 and Zm00001d024414), and a late embryogenesis abundant gene *ZmRAB28* (*responsive to abscisic acid28*, Zm00001d027740) (Niogret et al., 1996). We found that several of the MADS-box (named for yeast *minichromosomal maintenance* [*MCM1*], plant *AGAMOUS* [*AG*] and *DEFICIENS* [*DEF*], and human *serum response factor* [*SRF*]) TFs were also identified as seed-related signature genes, and may be involved in ovule development. Some such representative examples were *ZmMADS1* (Zm00001d023955), *ZmMADS6* (Zm00001d017614), *ZmMADS27* (Zm00001d006094), *ZmMADSL6* (Zm00001d031620), *ZmMADS24* (Zm00001d034047), and *ZmMADS7-LIKE* (Zm00001d021057).

Further analysis of these seed-related signature genes, together with the ChIP-Seq data of *O2* assayed at the stage of 15 DAP (Li et al., 2015), identified 32 signature genes as O2-modulated and/or -bound target genes (**Figure 3E**; **Table S7**), including α, β, and γ-zein genes (e.g., Zm00001d048809, Zm00001d048812, Zm00001d035760), as well as functionally unannotated genes (e.g., Zm00001d019925, Zm00001d020498, Zm00001d048810, and Zm00001d019156). Considering the characteristics of tissue specificity, these identified regulatory relationships between seed-signature genes would be valuable for investigating regulatory mechanisms occurring specifically in maize seed.

### Use of easyMF to reveal the relationship between time after pollination and gene expression during early maize seed development

Of 285 seed samples, 62 were harvested for 31 time points (two biological replicates per time point) from four stages of early maize seed development: at about double fertilization (0~16 hours after pollination [HAP]; stage I), coenocyte formation (20~44 HAP; stage II), cellularization (48~102 HAP; stage III), and differentiation (108~144 HAP; stage IV) (Yi et al., 2019). Using these temporal transcriptomes, a gene expression matrix G_t_ of 22,428 protein-coding genes and 31 time points was constructed in which each gene had FPKM ≥ 1 in at least one time point. In the following, we illustrate the application of easyMF to Gt to explore the temporal effect on gene expression during early maize seed development.

The NMF algorithm with easyMF was used to decompose Gt into a product of two matrices, namely amplitude matrix AM_t_ and pattern matrix PM_t_ (**Figure 4A**). Hierarchical clustering of AMt showed that 31 time points can be grouped into three sets, corresponding to three metagenes: metagene1 for stages I and II of the maize seed development, metagene2 for stage III, and metagene3 for stage IV. The easyMF platform grouped time points from stages I and II into one set, consistent with the hierarchical clustering analysis of the gene expression matrix Gt, where samples from the stages I and II were clustered in the same main branch (Yi et al., 2019). In our work, easyMF identified 1,250, 698, and 1,219 signature genes with peak expressions at stages I+II, III, and IV, respectively (**Table S7**). Three representative examples, including *ZmUMC1966* (*Zm00001d016705*) for metagene1, *ZmZNOD1* (*Zea nodulation homolog1*; Zm00001d045302) for metagene2, and *ZmFL3* (*floury3*; Zm00001d009292) for metagene3, are shown in **Figure 4B**.

**Figure 4.**
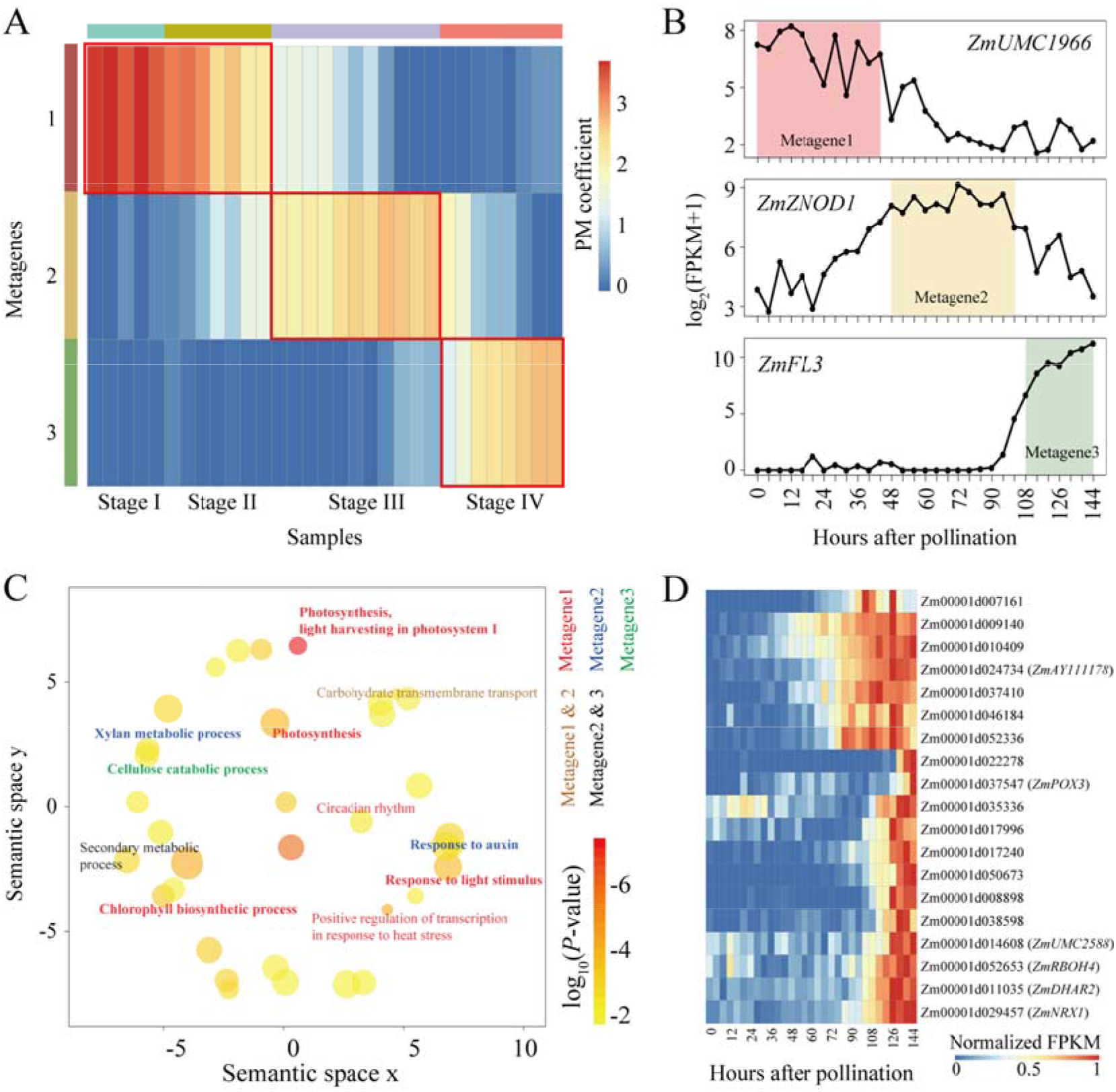
Application of easyMF to temporal transcriptome analysis. (A) Heatmap of the pattern matrix PM_t_ decomposed from temporal transcriptomes of early developmental stages of maize seed. (B) Expression profiles of *ZmUMC1966* (*Zm00001d016705*), *ZmZNOD1* (*Zea nodulation homolog1*; Zm00001d045302), and *ZmFL3* (*floury3*; Zm00001d009292). (C) GO enrichment results of signature genes from three metagenes. (D) Heatmap showing expression levels of 19 genes with annotations enriched with the “cellular oxidant detoxification” term.

For each set of signature genes, easyMF implemented topGO (Alexa and Rahnenführer, 2009) to perform GO enrichment analysis for the purpose of identifying important biological processes involved in the maize early seed development. This exploratory analysis revealed the importance of photosynthesis at approximately the double fertilization and coenocyte formation stages. Several signature genes from metagene1, including *ZmPSB29* (*photosystem II subunit29*, Zm00001d021763), *ZmPSA2* (*photosystemI2*, Zm00001d031738), and three oxygen-evolving complex genes (Zm00001d036535, Zm00001d021703, Zm00001d014564) (Pal et al., 2013; Vogt et al., 2015) are enriched in the photosynthesis system, corresponding to GO terms such as “photosynthesis, light harvesting in photosystem I”, “response to light stimulus”, and “photosystem II assembly” (**Figure 4C** and **Table S8**). Auxin has been reported to regulate cell fate specification at cellularization (Pagnussat et al., 2009), and xylose has been reported to be the most abundant monosaccharide constituent of maize cell walls (Jung and Casler, 2006). Several cell wall-related genes that may play roles during the cellularization stage were identified by using easyMF, including two cell wall invertase-related genes *incw1 (cell wall invertase 1*; Zm00001d016708) and *incw5* (*cell wall invertase 5*; Zm00001d025354). Twenty-three signature genes from metagene2 were also identified to respond to auxin and to be involved in the xylan metabolic process (**Table S8**). Cellular oxidant detoxification was linked according to the easyMF analysis with the initial endosperm differentiation stage. From metagene3, 19 signature genes, including *ZmDHAR2* (Zm00001d011035), *ZmUMC2588* (Zm00001d014608), *ZmNRX1* (Zm00001d029457), *ZmPOX3* (Zm00001d037547), and *ZmRBOH4* (Zm00001d052653), were indicated to possibly participate in cellular oxidant detoxification (**Table S8**), and to be expressed more abundantly at the initial endosperm differentiation stage than at the cellularization phase (**Figure 4D**).

In summary, these results indicated the value of using easyMF to extract expression patterns from temporal transcriptomes for the purpose of determining the responses of signature genes to developmental time, and consequently its value also in contributing to gaining a better understanding of the biology of specific developmental phases.

### Use of easyMF to attain compartment-specific biological knowledge from maize seed spatial transcriptomes

Finally, we focused on the spatial transcriptomes profiled from 10 compartments of maize kernel at 8 DAP (**Figure 5A**), with these 10 compartments including the aleurone (AL), the basal endosperm transfer layer (BETL), the embryo-surrounding region (ESR), the central starchy endosperm (CSE), the conducting zone (CZ), the embryo (EMB), the nucellus (NU), the placento-chalazal region (PC), the pericarp (PE), and the vascular region of the pedicel (PED) (Zhan et al., 2015). Using these transcriptomes, a gene expression matrix G_s_ (22,998 genes × 10 compartments) was constructed, in which each gene had FPKM ≥ 1 in at least one compartment. The easyMF platform was then tested for its usefulness in the analysis of spatial transcriptomes, specifically by decomposing the expression matrix Gs with the NMF algorithm and varying the number of metagenes.

**Figure 5.**
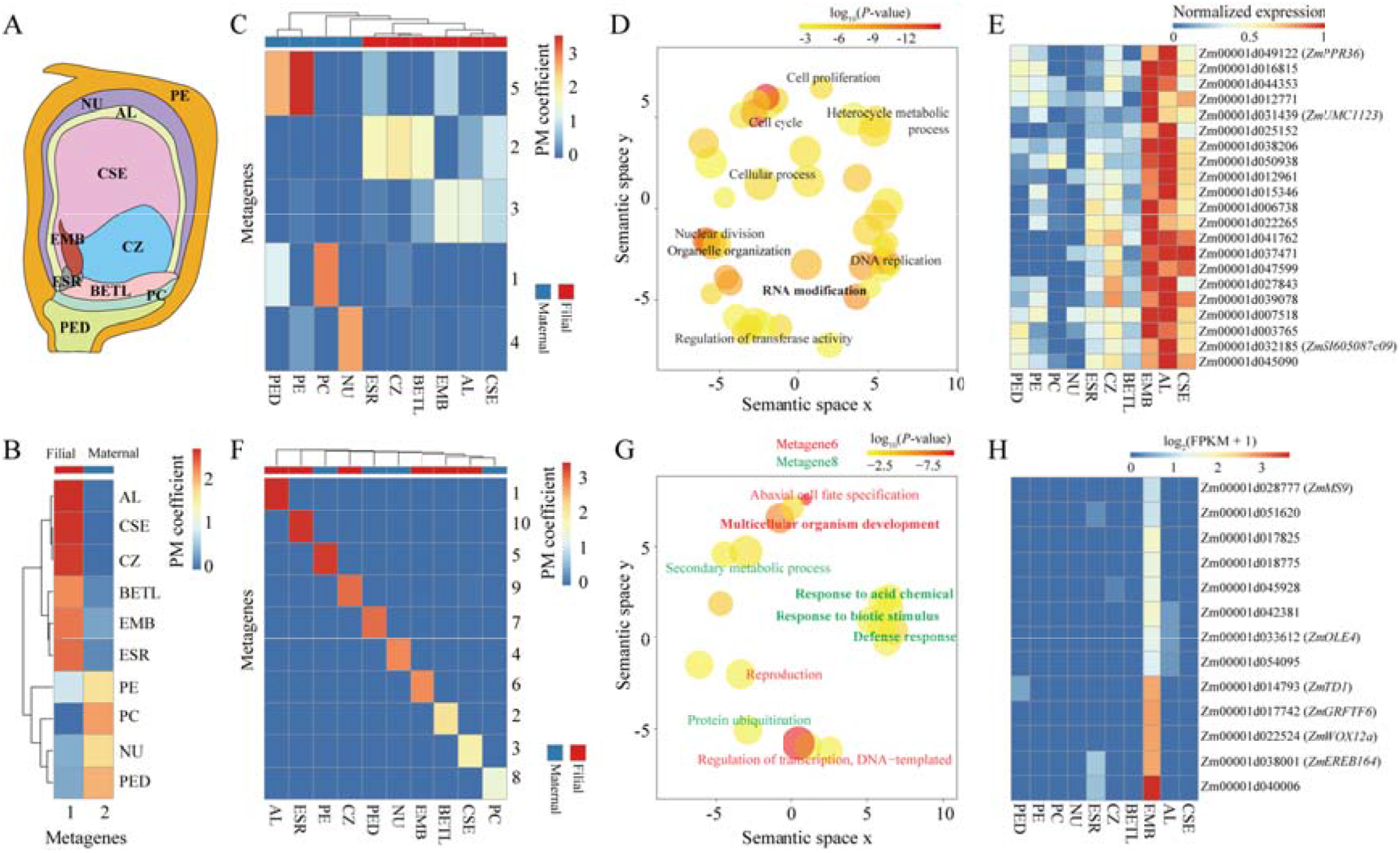
Application of easyMF to spatial transcriptome analysis. (A) Scheme representing 10 compartments of maize kernel at 8 DAP. (B) Hierarchical clustering analysis of the pattern matrix grouped 10 compartment samples from filial and maternal tissues into two distinct classes when the number of metagenes was set to two. (C) Hierarchical clustering analysis of the pattern matrix grouped spatially adjacent compartments into the same metagene when the number of metagenes was set to five. (D) GO enrichment results of signature genes from the metagene related to EMB, AL, and CSE compartments. (E) Heatmap exhibiting expression levels of 21 signature genes with annotations enriched with the “RNA modification” term. (F) Hierarchical clustering analysis of the pattern matrix identified compartment-specific metagenes when the number of metagenes was set to ten. (G) GO enrichment results of signature genes from EMB- and PC-related metagenes. (H) Heatmap exhibiting expression levels of 13 signature genes with annotations enriched with the “post-embryonic development” term.

We found that easyMF can distinguish compartments from filial and maternal tissues of maize kernel when setting the number of metagenes to be two. Hierarchical clustering of the generated pattern matrix showed that compartments belonging to filial and maternal tissues of 8-DAP maize kernel were perfectly grouped into two metagenes, respectively (**Figure 5B**). When increasing the number of metagenes, easyMF was able to identify signature genes and biological processes associated with specific maize kernel compartments. For example, when the number of metagenes was set to five, three spatially adjacent compartments were assigned to the same metagene (specifically, EMB, AL, and CSE for metagene3) (**Figure 5C**). Use of easyMF led to the identification of 286 signature genes for metagene3 (**Table S7**), GO analysis of which revealed a significant enrichment in GO terms related to DNA replication, cell cycle, nuclear division, and organelle organization (**Figure 5D**; **Table S9**), consistent with the extensive developmental and mitotic activity within these three compartments at this stage (Doll et al., 2020). Interestingly, the GO term “RNA modification” was also found to be significantly enriched, and was found in the annotations of 21 signature genes including 15 genes encoding pentatricopeptide repeat-containing proteins such as Zm00001d012961, Zm00001d015346, and Zm00001d016815, with high expression levels in the EMB, AL, and CSE compartments (**Figure 5E**). When the number of metagenes was increased to ten, easyMF extracted a different spatial expression pattern for each of these 10 compartments (**Figure 5F**), allowing for the identification of compartment-specific genes and different biological processes. For example, in the EMB-related metagene (i.e., metagene6), several signature genes such as *ZmWOX12a* (wuschel-related homeobox12A, Zm00001d022524) (Wu et al., 2007) and *ZmOLE4* (oleosin4, Zm00001d033612) (Miquel et al., 2014) were associated with organism development, corresponding to the GO terms “post-embryonic development”, “cell differentiation”, and “reproductive system development” (**Figure 5G-H; Table S7; Table S9**). In contrast, several signature genes (e.g., *ZmIAA40* (Aux/IAA-transcription factor 40, Zm00001d044818), *ZmPIN12* (PIN-formed protein12, Zm00001d045219), and *ZmZIM14* (ZIM-transcription factor 14, Zm00001d048268)) from the metagene8 (PC compartment) were enriched in terms indicating processes involving response to stimulus such as “defense response”, “response to abscisic acid”, and “response to biotic stimulus” (**Figure 5G**; **Table S7; Table S9**).

Taken together, as a result of an MF-based analysis of maize seed spatial transcriptomes, it was shown that easyMF could be used to open a window for attaining compartment-specific biological knowledge with the discovery of signature genes and related biological processes, and its use here specifically enhanced our understanding of the process of cell differentiation during seed development.

## Discussion

With the ever-increasing volumes of RNA-Seq data, the use of MF has been a foundational approach to extracting biological knowledge in transcriptomic studies. Although a variety of MF-based software packages are already available (**Table S1**), many of them have limitations, including not being easy to use, and not providing an all-in-one solution in the transcriptome data analysis. We compared the features provided in the easyMF platform with those in available MF-based software packages, and describe here three major differences.

One distinct difference involves their capabilities of preparing a high-quality gene expression matrix, which is essential for knowledge discovery through MF-based and other forms of transcriptome analysis. All currently available MF-based software packages only accept a gene expression matrix as input, and do not provide for the option to process raw RNA-Seq data to gene expression values (**Table S1**). This limitation hinders the ability to effectively analyze RNA-Seq data generated locally by the user or deposited in public repositories (e.g., the NCBI GEO and SRA databases). In addition, they lack a quality control function that filters genes expressed at low levels and/or outlier samples for downstream analysis. The easyMF platform was designed to address these two limitations by incorporating a customized RNA-Seq analysis pipeline (**Figure S1**), which is invaluable for researchers with relatively little experience in high-throughput RNA-Seq data analysis.

Another notable difference involves the comprehensiveness of the MF-based analysis. Almost all currently available MF tools, except Compadre (Ramos-Rodriguez et al., 2012), were designed with a focus on only one of the three above-mentioned MF algorithms (PCA, ICA, and NMF) and have limited embedded functionalities for metagene-based exploratory analysis (**Table S1**). In contrast, easyMF was specially designed to implement all three of these MF algorithms with R packages: ‘stats’ (*prcomp*) for PCA (Team, 2018), ‘ica’ (*icafast*) for ICA (Helwig, 2018), and ‘bignmf’ (*bignmf*) for NMF (Pan et al., 2012). It can be used to perform a series of metagene-based exploratory analytical tasks through sample clustering, signature gene identification, functional gene discovery, and pathway activity inference (**Figure 1**). These features allow easyMF to serve as an all-in-one platform for comprehensively mining large-scale gene expression data using MF algorithms.

The third and last major difference involves the flexibility of use. Most currently available MF tools were produced as bioinformatics toolkits with command-based implementations, and lack intuitive representations of the results of the analyses. But easyMF was developed as a Galaxy-based platform that aims to easily perform MF-based analysis. Taking advantage of the Galaxy system, easyMF provides user-friendly GUIs to design bioinformatics pipelines with different functionalities, handle large volumes of RNA-Seq data, adjust different input parameters, examine the running status, and visualize output results. It also permanently records all analysis data such as inputs, parameters, intermediate results, and outputs in the ‘history’ panel of the easyMF platform, making complex MF-based transcriptomic analysis reproducible and amenable to collaborative modes. Moreover, easyMF is packaged into a Docker image that can be employed under different operating systems (i.e., Windows, Linux, and Macintosh), overcoming issues related to code changes, library dependencies and backward compatibility over time. We expect the easy implementation of easyMF as well as the detailed user documentation and open-access wiki discussion groups to allow researchers, regardless of their levels of programming experience, to carry out accessible, reproducible and collaborative analyses of large-volume of RNA-Seq data with MF algorithms.

We have demonstrated the use of easyMF in the MF-based analysis of maize transcriptomes for four case studies: (1) prioritization of seed-related genes, (2) clustering analysis of seed samples, (3) temporal analysis of maize seed transcriptomes, and (4) spatial analysis of maize seed transcriptomes. The new knowledge attained about maize seeds using easyMF illustrated that this tool is readily applicable and flexible to confront a range of biological questions, allowing users to more effectively concentrate on hypothesis testing. There were also some limitations regarding the study. First, the efficiency of easyMF was illustrated only using 940 RNA-Seq datasets from maize. The ability of easyMF to handle more transcriptome data and more complex species (e.g., hexaploid bread wheat) needs to be investigated in future work. Secondly, we did not show the application of easyMF in the analysis of single-cell transcriptome data (**File S1; Figure S5**), which were deficient for maize kernels at the time the work was carried out. Single-cell RNA-Seq, which measures gene expressions at the level of a single cell, has been developed as a powerful method to investigate the function of individual cells (Tang et al., 2010). Using single-cell RNA-Seq data from *Arabidopsis* root (Shulse et al., 2018), we found that easyMF can be used to map six cell types according to 13 clusters of 4,043 cells (**Figure S6**). Thirdly and lastly, large-scale transcriptome analysis using MF and other algorithms is often time-consuming and requires heavy computational resources. Despite the easy deployment and implementation of easyMF, it still cannot be used by researchers lacking high-throughput computational resources. In such cases, we would happily collaborate on analyses, and such a collaboration can be requested by contacting the corresponding author.

The easyMF project is still being developed and improved. The source codes, user manual, Docker image, prototype web server and all future updates are available at the homepage of easyMF project (https://github.com/cma2015/easyMF). The easyMF Docker image can be obtained at https://hub.docker.com/r/malab/easymf. A porotype web server for easyMF has been developed by the Aliyun cloud computing architecture and can be accessed at http://easymf.omicstudio.cloud.

## Methods

### Generation of the maize gene expression matrix G_1_

easyMF presents a customized bioinformatics pipeline to generate gene expression matrix from raw RNA-Seq reads (**Figure S1**). This bioinformatics pipeline has been applied to generate the maize gene expression matrix G_1_. In brief, 1,066 maize RNA-Seq datasets were firstly collected from NCBI’s Gene Expression Omnibus (GEO) and/or Sequence Read Archive (SRA) databases (as of 26 July 2019). Raw RNA-Seq data were preprocessed using fastp (version 0.20.0) (Chen et al., 2018) for quality control, including sequencing adapter trimming and low-quality read filtering. Subsequently, high-quality RNA-Seq reads from each sample were aligned to maize reference genome (APGv4, https://plants.ensembl.org/Zea_mays/Info/Index) using HISAT2 (version 2.1.0) (Kim et al., 2015), generating a BAM (binary alignment map) file recording read-genome alignments. BAM files were then used as inputs of StringTie (version 1.3.6) (Pertea et al., 2015) to estimate gene expression abundance. To obtain a high-quality gene expression matrix, a two-step quality control was implemented to filter genes expressed at low levels and remove outlier samples. For expression-level quality control, genes with FPKM (fragments per kilobase of transcript per million mapped reads) ≥ 1 in at least 15 RNA-Seq samples were retained. For low-quality samples, we firstly averaged the statistical duplicated samples based on PCC with criteria: PCC > 0.999 (Fehrmann et al., 2015). Then, outlier samples whose correlation with the first principal component (sample based principal component analysis) less than 0.75 were removed. Finally, a gene expression matrix G_1_ with 28,874 protein-coding genes and 940 samples was obtained for the downstream application.

### Identification of signature genes

easyMF decomposes a high-dimensional gene expression matrix (genes in rows and samples in columns) into a product of two low-dimensional metagene-based matrices: an amplitude matrix (AM; genes in rows and metagenes in columns) and a pattern matrix (PM; metagenes in rows and samples in columns). Using gene-level relationships in the AM and sample-level relationships in the PM, easyMF identifies genes exhibiting dominant patterns (defined as signature genes) for each metagene using the patternMarkers (Stein-O’Brien et al., 2017) and the Pearson’s correlation coefficient (PCC) statistics. patternMarkers calculates a Euclidean distance (*ED*) between normalized AM coefficients (i.e., coefficients in the AM) and a 0-1 pattern of metagenes (**Figure S2A**). Suppose the number of metagenes is *M*, the *ED* score for gene *i* and metagene *k* (1≤ *k ≤ M*) is calculated using following formula:

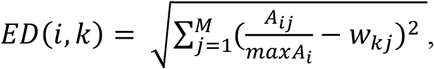

where *A_ij_* represents the AM coefficient of gene *i* in metagene *j* (1≤ *j ≤ M*), *maxA_i_*, denotes the maximum value of AM coefficient of gene *i* among all *M* metagenes, 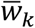 is a numeric unit vector specifying the status of each component (*w_kj_* = 1 only when *k* = *j*, otherwise equals to 0). For each gene *i*, easyMF repeats this process to generate a vector of *ED* scores for all metagenes.

easyMF uses the PCC statistic to quantify the correlation between gene expression abundance and PM coefficients (**Figure S2B**). For gene *i* and metagene *k*, the PCC score is calculated using the following formula:

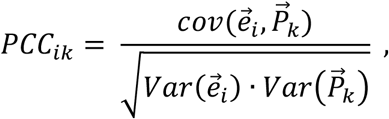

Where 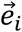 represents the *i*-th gene’s expression values, 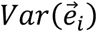 represents the variance of *i*-th gene’s expression values, 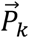 represents the *i*-th gene’s PM coefficients, 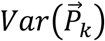 represents the variance of *i*-th gene’s PM coefficients. The gene *i* is regarded as a signature gene of metagene *k*, if it satisfied with three conditions: i) equals to the minimal ED score; ii) PCC ≥ 0.6; iii) *P*-value ≤ 1.0E-03. Of note, the thresholds of PCC and *P*-value can be user-adjusted in web interface of easyMF.

### Metagene-based gene prioritization

For a given set of genes (denoted as labeled genes), easyMF firstly examines the difference in the distribution of AM coefficients between labeled genes and unlabeled (all except labeled genes in the AM) genes by using Student’s t-test, following by a transformation of the significance level *P*-value to *z*-score using the standard normal quantile function ‘qnorm’ in R. A higher *z*-score indicates a larger difference in the AM coefficient between labeled and unlabeled genes, thus corresponding to stronger biological association between the metagene and the gene set. This results to a *z*-score vector with a length of metagene number for the given gene set. Subsequently, the association between *z*-scores and AM coefficients of corresponding genes is examined using the PCC statistic (Fehrmann et al., 2015). Finally, easyMF prioritizes candidate genes functionally associated with the given gene set based on the decreasing PCC values (**Figure S3**).

### Performance evaluation of gene prioritization approaches

The leave-one-out cross-validation (LOOCV) experiment was used to evaluate the performance of easyMF and MaizeNet (Lee et al., 2019) in gene prioritization. In LOOCV experiment, each labeled gene and all unlabeled genes were used as testing samples and their normalized scores (min-max normalization) were then calculated. The performance of easyMF and MaizeNet were further evaluated using the area under the receiver operating characteristic (ROC) curve (AUC) and the area under the curve of self-ranked curve (AUSR). The ROC curve is a plot of true-positive rate (TPR) along the y axis versus false-positive rate (FPR) along the x axis. While the self-rank curve is a plot of ratio (*Ra*) along the *y* axis versus self-rank along the *x* axis (Tzfadia et al., 2012), the *Ra* can be calculated by the following formula:

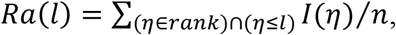

Where *rank* represents the ranks of all positive genes, *Ra*(*T*) represents the ratio of ranks lower than a pre-defined level of *l* (e.g., 1000). Both AUC and AUSR, ranging from 0 to 1, were finally calculated using the trapezoid rule (Radivojac et al., 2013), with greater value indicating better prediction performance.

## Supplementary Data

**All Supplemental tables (Table S1-9) are available at the Zenodo Public Data Repository (http://doi.org/10.5281/zenodo.4383238).**

**File S1.** Metagene-based pathway activity analysis and single-cell RNA-Seq data analysis.

**Table S1**. Summary of MF-based software tools.

**Table S2**. Description of functional modules in easyMF.

**Table S 3**. Summary of 940 maize RNA-Seq datasets used in this study.

**Table S4**. Summary of 75 GO terms used to evaluate the performance of gene prioritization methods.

**Table S5**. List of 70 experimentally validated genes functionalized in maize seed development.

**Table S6**. Genome-wide prioritization of candidate seed-related genes using easyMF and MaizeNet.

**Table S7**. List of signature genes identified from gene expression matrix G_1_, G_t_, and Gs.

**Table S8**. GO enrichments of signature genes identified from gene expression matrix Gt.

**Table S9**. GO enrichments of signature genes identified from gene expression matrix Gs.

**Figure S1.** The bioinformatics pipeline for the generation of a gene expression matrix from RNA-Seq data.

**Figure S2**. Identification of signature genes using patternMarkers (A) and Pearson’s correlation coefficient (PCC) algorithm (B).

**Figure S3.** Prioritization of candidate genes involved in a pre-specific function.

**Figure S4**. PCA statistics of optimal metagenes. The blue and black dots represent the Cronbach’s α value and the explained variance of each metagene, respectively. The red dots represent the cumulative explained variance. At the threshold of Cronbach’s *α* of 0.7, easyMF generated 161 optimal metagenes, capturing 96.4% of the variation in gene expression.

**Figure S5**. Identification of cell types of unknown cells from single-cell RNA-Seq data.

**Figure S6**. t-Distributed Stochastic Neighbor Embedding (t-SNE) dimensionality reduction of 4,043 single *Arabidopsis* root cells, which are represented by individual points. All captured cells were clustered into 13 populations corresponding to six cell types.

## Acknowledgements

We thank High-Performance Computing (HPC) of Northwest A&F University for providing computing resources. This work was supported by the National Natural Science Foundation of China (31570371), the Youth 1000-Talent Program of China, the Hundred Talents Program of Shaanxi Province of China, Projects of Youth Technology New Star of Shaanxi Province (2017KJXX-67), and the Fundamental Research Funds for the Central Universities (2452020041).

## Author Contributions

C.M. conceived the project; W.M. and S.C. developed the platform and performed the applications; W.M., S.C., J.Z., S.X., and M.S. tested the platform. S.C., W.M., Y.Q. and C.M. explained the application results; S.C., W.M., and C.M. wrote the article. All authors read, revised, and approved the final manuscript.

## References

Abdi, H., and Williams, L.J. (2010). Principal component analysis. Wiley Interdisciplinary Reviews: Computational Statistics 2, 433–459.

Alexa, A., and Rahnenführer, J. (2009). Gene set enrichment analysis with topGO. https://bioconductor.riken.jp/packages/3.2/bioc/vignettes/topGO/inst/doc/topGO.pdf.

Bernardi, J., Battaglia, R., Bagnaresi, P., Lucini, L., and Marocco, A. (2019). Transcriptomic and metabolomic analysis of *ZmYUC1* mutant reveals the role of auxin during early endosperm formation in maize. Plant Sci 281, 133–145.

Bernardi, J., Lanubile, A., Li, Q.B., Kumar, D., Kladnik, A., Cook, S.D., Ross, J.J., Marocco, A., and Chourey, P.S. (2012). Impaired auxin biosynthesis in the defective endosperm18 mutant is due to mutational loss of expression in the *ZmYUC1* gene encoding endosperm-specific YUCCA1 protein in maize. Plant Physiol 160, 1318–1328.

Bodenhofer, U., Kothmeier, A., and Hochreiter, S. (2011). APCluster: an R package for affinity propagation clustering. Bioinformatics 27, 2463–2464.

Bolser, D.M., Staines, D.M., Perry, E., and Kersey, P.J. (2017). Ensembl Plants: integrating tools for visualizing, mining, and analyzing plant genomic data. Methods Mol Biol 1533, 1–31.

Cardoso-Moreira, M., Halbert, J., Valloton, D., Velten, B., Chen, C., Shao, Y., Liechti, A., Ascencao, K., Rummel, C., Ovchinnikova, S., Mazin, P.V., Xenarios, I., Harshman, K., Mort, M., Cooper, D.N., Sandi, C., Soares, M.J., Ferreira, P.G., Afonso, S., Carneiro, M., Turner, J.M.A., VandeBerg, J.L., Fallahshahroudi, A., Jensen, P., Behr, R., Lisgo, S., Lindsay, S., Khaitovich, P., Huber, W., Baker, J., Anders, S., Zhang, Y.E., and Kaessmann, H. (2019). Gene expression across mammalian organ development. Nature 571, 505–509.

Chen, S., Zhou, Y., Chen, Y., and Gu, J. (2018). fastp: an ultra-fast all-in-one FASTQ preprocessor. Bioinformatics 34, i884–i890.

Cuocolo, R., Perillo, T., De Rosa, E., Ugga, L., and Petretta, M. (2019). Current applications of big data and machine learning in cardiology. J Geriatr Cardiol 16, 601–607.

Dixon, P.J.J.o.V.S. (2003). VEGAN, a package of R functions for community ecology 14, 927–930.

Doll, N.M., Just, J., Brunaud, V., Caius, J., Grimault, A., Depege-Fargeix, N., Esteban, E., Pasha, A., Provart, N.J., Ingram, G.C., Rogowsky, P.M., and Widiez, T. (2020). Transcriptomics at maize embryo/endosperm interfaces identifies a transcriptionally distinct endosperm subdomain adjacent to the embryo scutellum. Plant Cell 32, 833–852.

Fehrmann, R.S., Karjalainen, J.M., Krajewska, M., Westra, H.J., Maloney, D., Simeonov, A., Pers, T.H., Hirschhorn, J.N., Jansen, R.C., Schultes, E.A., van Haagen, H.H., de Vries, E.G., te Meerman, G.J., Wijmenga, C., van Vugt, M.A., and Franke, L. (2015). Gene expression analysis identifies global gene dosage sensitivity in cancer. Nature Genetics 47, 115–125.

Feng, F., Qi, W., Lv, Y., Yan, S., Xu, L., Yang, W., Yuan, Y., Chen, Y., Zhao, H., and Song, R. (2018). OPAQUE11 is a central hub of the regulatory network for maize endosperm development and nutrient metabolism. Plant Cell 30, 375–396.

Flint-Garcia, S.A., Bodnar, A.L., and Scott, M.P. (2009). Wide variability in kernel composition, seed characteristics, and zein profiles among diverse maize inbreds, landraces, and teosinte. Theor Appl Genet 119, 1129–1142.

Gaujoux, R., and Seoighe, C. (2010). A flexible R package for nonnegative matrix factorization. BMC Bioinformatics 11, 367.

Guo, M., Rupe, M.A., Danilevskaya, O.N., Yang, X., and Hu, Z. (2003). Genome-wide mRNA profiling reveals heterochronic allelic variation and a new imprinted gene in hybrid maize endosperm. Plant J 36, 30–44.

Haun, W.J., and Springer, N.M. (2008). Maternal and paternal alleles exhibit differential histone methylation and acetylation at maize imprinted genes. Plant J 56, 903–912.

Helwig, N.E. (2018). ica: independent component analysis. http://search.r-project.org/library/ica/html/ica-package.html

Hennig, C. (2013). fpc: flexible procedures for clustering. https://cran.r-project.org/web/packages/fpc/index.html.

Hyvarinen, A., and Oja, E. (2000). Independent component analysis: algorithms and applications. Neural Networks 13, 411–430.

Jung, H.G., and Casler, M.D. (2006). Maize stem tissues. Crop Science 46, 1801.

Kim, D., Langmead, B., and Salzberg, S.L. (2015). HISAT: a fast spliced aligner with low memory requirements. Nat Methods 12, 357–360.

Koren, Y., Bell, R., and Volinsky, C. (2009). Matrix factorization techniques for recommender systems. Computer 42, 30–37.

Kryuchkova-Mostacci, N., and Robinson-Rechavi, M. (2017). A benchmark of gene expression tissue-specificity metrics. Brief Bioinform 18, 205–214.

Lee, D., and Seung, H.S. (2000). Algorithms for non-negative matrix factorization. Advances in neural information processing systems 13, 556–562.

Lee, T., Lee, S., Yang, S., and Lee, I. (2019). MaizeNet: a co-functional network for network-assisted systems genetics in *Zea mays*. Plant J 99, 571–582.

Leek, J.T., Johnson, W.E., Parker, H.S., Jaffe, A.E., and Storey, J.D. (2012). The sva package for removing batch effects and other unwanted variation in high-throughput experiments. Bioinformatics 28, 882–883.

Li, C., Qiao, Z., Qi, W., Wang, Q., Yuan, Y., Yang, X., Tang, Y., Mei, B., Lv, Y., Zhao, H., Xiao, H., and Song, R. (2015). Genome-wide characterization of *cis*-acting DNA targets reveals the transcriptional regulatory framework of *opaque2* in maize. Plant Cell 27, 532–545.

Lopez, M., Gomez, E., Faye, C., Gerentes, D., Paul, W., Royo, J., Hueros, G., and Muniz, L.M. (2017). ZmSBT1 and *ZmSBT2,* two new subtilisin-like serine proteases genes expressed in early maize kernel development. Planta 245, 409–424.

Ma, C., Zhang, H.H., and Wang, X. (2014). Machine learning for Big Data analytics in plants. Trends Plant Sci 19, 798–808.

Ma, C., Li, B., Wang, L., Xu, M.L., Lizhu, E., Jin, H., Wang, Z., and Ye, J.R. (2019). Characterization of phytohormone and transcriptome reprogramming profiles during maize early kernel development. BMC Plant Biol 19, 197.

Maechler, M., Rousseeuw, P., Struyf, A., Hubert, M., and Hornik, K. (2012). Cluster: cluster analysis basics and extensions 1, 56.

Miquel, M., Trigui, G., d’Andrea, S., Kelemen, Z., Baud, S., Berger, A., Deruyffelaere, C., Trubuil, A., Lepiniec, L., and Dubreucq, B. (2014). Specialization of oleosins in oil body dynamics during seed development in *Arabidopsis* seeds. Plant Physiol 164, 1866–1878.

Mooney, S.J., and Pejaver, V. (2018). Big Data in public health: terminology, machine learning, and privacy. Annu Rev Public Health 39, 95–112.

Nelms, B., and Walbot, V. (2019). Defining the developmental program leading to meiosis in maize. Science 364, 52–56.

Nguyen, N.D., and Wang, D. (2020). Multiview learning for understanding functional multiomics. PLoS Comput Biol 16, e1007677.

Niogret, M.F., Culianez-Macia, F.A., Goday, A., Mar Alba, M., and Pages, M. (1996). Expression and cellular localization of *rab28* mRNA and Rab28 protein during maize embryogenesis. Plant J 9, 549–557.

Noor, E., Cherkaoui, S., and Sauer, U. (2019). Biological insights through omics data integration. Current Opinion in Systems Biology 15, 39–47.

One Thousand Plant Transcriptomes, I. (2019). One thousand plant transcriptomes and the phylogenomics of green plants. Nature 574, 679–685.

Pagnussat, G.C., Alandete-Saez, M., Bowman, J.L., and Sundaresan, V. (2009). Auxin-dependent patterning and gamete specification in the *Arabidopsis* female gametophyte. Science 324, 1684–1689.

Pal, R., Negre, C.F., Vogt, L., Pokhrel, R., Ertem, M.Z., Brudvig, G.W., and Batista, V.S. (2013). S0-State model of the oxygen-evolving complex of photosystem II. Biochemistry 52, 7703–7706.

Pan, L., Qiu, Y., and Wei, T. (2012). bignmf: Solving NMF via coordinate descent. https://github.com/panlanfeng/bignmf.

Pertea, M., Pertea, G.M., Antonescu, C.M., Chang, T.C., Mendell, J.T., and Salzberg, S.L. (2015). StringTie enables improved reconstruction of a transcriptome from RNA-seq reads. Nat Biotechnol 33, 290–295.

Qiu, Z., Chen, S., Qi, Y., Liu, C., Zhai, J., Xie, S., and Ma, C. (2020). Exploring transcriptional switches from pairwise, temporal and population RNA-Seq data using deepTS. Brief Bioinform. doi: 10.1093/bib/bbaa137.

Radivojac, P., Clark, W.T., Oron, T.R., Schnoes, A.M., Wittkop, T., Sokolov, A., Graim, K., Funk, C., Verspoor, K., Ben-Hur, A., Pandey, G., Yunes, J.M., Talwalkar, A.S., Repo, S., Souza, M.L., Piovesan, D., Casadio, R., Wang, Z., Cheng, J.L., Fang, H., Goughl, J., Koskinen, P., Toronen, P., Nokso-Koivisto, J., Holm, L., Cozzetto, D., Buchan, D.W.A., Bryson, K., Jones, D.T., Limaye, B., Inamdar, H., Datta, A., Manjari, S.K., Joshi, R., Chitale, M., Kihara, D., Lisewski, A.M., Erdin, S., Venner, E., Lichtarge, O., Rentzsch, R., Yang, H.X., Romero, A.E., Bhat, P., Paccanaro, A., Hamp, T., Kassner, R., Seemayer, S., Vicedo, E., Schaefer, C., Achten, D., Auer, F., Boehm, A., Braun, T., Hecht, M., Heron, M., Honigschmid, P., Hopf, T.A., Kaufmann, S., Kiening, M., Krompass, D., Landerer, C., Mahlich, Y., Roos, M., Bjorne, J., Salakoski, T., Wong, A., Shatkay, H., Gatzmann, F., Sommer, I., Wass, M.N., Sternberg, M.J.E., Skunca, N., Supek, F., Bosnjak, M., Panov, P., Dzeroski, S., Smuc, T., Kourmpetis, Y.A.I., van Dijk, A.D.J., ter Braak, C.J.F., Zhou, Y.P., Gong, Q.T., Dong, X.R., Tian, W.D., Falda, M., Fontana, P., Lavezzo, E., Di Camillo, B., Toppo, S., Lan, L., Djuric, N., Guo, Y.H., Vucetic, S., Bairoch, A., Linial, M., Babbitt, P.C., Brenner, S.E., Orengo, C., Rost, B., Mooney, S.D., and Friedberg, I. (2013). A large-scale evaluation of computational protein function prediction. Nature Methods 10, 221–227.

Ramos-Rodriguez, R.R., Cuevas-Diaz-Duran, R., Falciani, F., Tamez-Pena, J.G., and Trevino, V. (2012). COMPADRE: an R and web resource for pathway activity analysis by component decompositions. Bioinformatics 28, 2701–2702.

Sarropoulos, I., Marin, R., Cardoso-Moreira, M., and Kaessmann, H. (2019). Developmental dynamics of lncRNAs across mammalian organs and species. Nature 571, 510–514.

Schmidt, R.J., Burr, F.A., Aukerman, M.J., and Burr, B. (1990). Maize regulatory gene *opaque-2* encodes a protein with a “leucine-zipper” motif that binds to zein DNA. Proc Natl Acad Sci U S A 87, 46–50.

Schmidt, R.J., Veit, B., Mandel, M.A., Mena, M., Hake, S., and Yanofsky, M.F. (1993). Identification and molecular characterization of *ZAG1,* the maize homolog of the *Arabidopsis* floral homeotic gene *AGAMOUS*. Plant Cell 5, 729–737.

Scrucca, L., Fop, M., Murphy, T.B., and Raftery, A.E. (2016). mclust 5: clustering, classification and density estimation using gaussian finite mixture models. R J 8, 289–317.

Shannon, J.C., Pien, F.M., Cao, H., and Liu, K.C. (1998). Brittle-1, an adenylate translocator, facilitates transfer of extraplastidial synthesized ADP--glucose into amyloplasts of maize endosperms. Plant Physiol 117, 1235–1252.

Shulse, C.N., Cole, B.J., Turco, G.M., Zhu, Y., Brady, S.M., and Dickel, D.E. (2018). High-throughput single-cell transcriptome profiling of plant cell types. bioRxiv 402966, doi: https://doi.org/10.1101/402966.

Sompairac, N., Nazarov, P.V., Czerwinska, U., Cantini, L., Biton, A., Molkenov, A., Zhumadilov, Z., Barillot, E., Radvanyi, F., Gorban, A., Kairov, U., and Zinovyev, A. (2019). Independent component analysis for unraveling the complexity of cancer omics datasets. Int J Mol Sci 20, 4414.

Stein-O’Brien, G.L., Arora, R., Culhane, A.C., Favorov, A.V., Garmire, L.X., Greene, C.S., Goff, L.A., Li, Y., Ngom, A., Ochs, M.F., Xu, Y., and Fertig, E.J. (2018). Enter the matrix: factorization uncovers knowledge from omics. Trends Genet 34, 790–805.

Stein-O’Brien, G.L., Carey, J.L., Lee, W.S., Considine, M., Favorov, A.V., Flam, E., Guo, T., Li, S., Marchionni, L., Sherman, T., Sivy, S., Gaykalova, D.A., McKay, R.D., Ochs, M.F., Colantuoni, C., and Fertig, E.J. (2017). PatternMarkers & GWCoGAPS for novel data-driven biomarkers via whole transcriptome NMF. Bioinformatics 33, 1892–1894.

Tang, F., Barbacioru, C., Nordman, E., Li, B., Xu, N., Bashkirov, V.I., Lao, K., and Surani, M.A. (2010). RNA-Seq analysis to capture the transcriptome landscape of a single cell. Nat Protoc 5, 516–535.

Team, R.C. (2018). R: A language and environment for statistical computing. https://www.r-project.org.

Tsai, C.Y. (1979). Tissue-specific zein synthesis in maize kernel. Biochem Genet 17, 1109–1119.

Tzfadia, O., Amar, D., Bradbury, L.M., Wurtzel, E.T., and Shamir, R. (2012). The MORPH algorithm: ranking candidate genes for membership in *Arabidopsis* and tomato pathways. Plant Cell 24, 4389–4406.

Vogt, L., Vinyard, D.J., Khan, S., and Brudvig, G.W. (2015). Oxygen-evolving complex of Photosystem II: an analysis of second-shell residues and hydrogen-bonding networks. Curr Opin Chem Biol 25, 152–158.

Wimalanathan, K., Friedberg, I., Andorf, C.M., and Lawrence-Dill, C.J. (2018). Maize GO annotation-methods, evaluation, and review (maize-GAMER). Plant Direct 2, e00052.

Wu, X., Chory, J., and Weigel, D. (2007). Combinations of *WOX* activities regulate tissue proliferation during *Arabidopsis* embryonic development. Dev Biol 309, 306–316.

Yi, F., Gu, W., Chen, J., Song, N., Gao, X., Zhang, X., Zhou, Y., Ma, X., Song, W., Zhao, H., Esteban, E., Pasha, A., Provart, N.J., and Lai, J. (2019). High temporal-resolution transcriptome landscape of early maize seed development. Plant Cell 31, 974–992.

Zhan, J., Li, G., Ryu, C.H., Ma, C., Zhang, S., Lloyd, A., Hunter, B.G., Larkins, B.A., Drews, G.N., Wang, X., and Yadegari, R. (2018). Opaque-2 regulates a complex gene network associated with cell differentiation and storage functions of maize endosperm. Plant Cell 30, 2425–2446.

Zhan, J., Thakare, D., Ma, C., Lloyd, A., Nixon, N.M., Arakaki, A.M., Burnett, W.J., Logan, K.O., Wang, D., Wang, X., Drews, G.N., and Yadegari, R. (2015). RNA sequencing of laser-capture microdissected compartments of the maize kernel identifies regulatory modules associated with endosperm cell differentiation. Plant Cell 27, 513–531.

Zhang, Z., Dong, J., Ji, C., Wu, Y., and Messing, J. (2019). NAC-type transcription factors regulate accumulation of starch and protein in maize seeds. Proc Natl Acad Sci U S A 116, 11223–11228.

